# Diisopropylfluorophosphate (DFP) volatizes and cross-contaminates wells in a common 96-well plate zebrafish larvae exposure method

**DOI:** 10.1101/2021.07.19.452994

**Authors:** Paige C. Mundy, Rosalia Mendieta, Pamela J. Lein

**Affiliations:** Department of Molecular Biosciences, University of California, School of Veterinary Medicine, Davis, CA 95616, United States of America

**Keywords:** Diisopropylfluorophosphate, organophosphate, zebrafish, toxicology

## Abstract

Diisopropylfluorophosphate (DFP) is an organophosphate (OP) that is commonly used to study the neurotoxic effects of acutely intoxicating OP exposure. In preliminary studies, we discovered abnormal deaths in DMSO-only exposed larvae housed in the same plate as DFP-exposed larvae, and hypothesized that DFP volatilizes and cross-contaminates wells when using a 96-well plate exposure method for exposing zebrafish larvae. Survivability and acetylcholinesterase activity assays confirmed DFP presence in the tissues of zebrafish ostensibly exposed to DMSO only. These findings indicate DFP cross-contamination, which raises concerns for the experimental design of studies evaluating the toxicity of volatile and semi-volatile substances.

## INTRODUCTION

Organophosphates (OPs) are a class of highly toxic compounds used extensively as insecticides (e.g., chlorpyrifos) around the world that are pervasive contaminants of diverse environmental compartments (Gunnell et al. 2007; DiBartolomeis et al. 2019). Some OPs (e.g., sarin, soman) have been weaponized as nerve agents, and self-poisonings with OP pesticides are estimated to occur at a rate of approximately 200,000 per year in developing countries (Eddleston et al. 2008). Non-fatal outcomes of acute OP poisonings can be severe and long-lasting, including onset of acquired epilepsy and persistent neurological deficits in humans (Chen 2012; Jett et al. 2020) and experimental rodent models (Hobson et al. 2019; González et al. 2020; Guignet et al. 2020). Therefore, acute OP intoxication is of worldwide importance. Because of the severity of exposure outcomes, there is great interest in characterizing and understanding the toxicological effects of OPs for setting exposure limits for agricultural runoff and drinking water, and identifying novel, mechanistically relevant therapeutic targets for mitigating adverse effects.

Diisopropylfluorophosphate (DFP) is often used to study acute OP intoxication in the laboratory (González et al. 2020; Guignet et al. 2020). The primary mechanism of action (MOA) of DFP and most other OPs is acetylcholinesterase (AChE) inhibition, which leads to the accumulation of acetylcholine in cholinergic synapses within the central and peripheral nervous systems (Hulse et al. 2014). Due to challenges of analyzing the highly reactive and short-lived DFP, AChE inhibition is often used as a surrogate marker that DFP has effectively penetrated target tissue and engaged its molecular target (González et al. 2020).

While rats have been extensively used as a preclinical model of acute DFP poisoning (Pessah et al. 2016; Pouliot et al. 2016), the neurotoxic effects of DFP have also been investigated in zebrafish, a common teleost model (Faria et al. 2018; Brenet et al. 2020).

Zebrafish are excellent models for medium to high-throughput toxicological assays due to their rapid generation rate and high fecundity (Cassar et al. 2020; Tal et al. 2020). Since they can survive in small amounts of aqueous media, exposing or evaluating zebrafish larvae in 96-well plates is an extensively used method in toxicology research and can be particularly powerful when screening large chemical libraries or phenotypically assessing mutant models (Dach et al. 2019; Colón-Rodríguez et al. 2020; Tal et al. 2020; Truong et al. 2020). A challenge is that compounds of interest need to be sufficiently water soluble for testing (Cassar et al. 2020; Tal et al. 2020). Microinjection is a possible alternative to aqueous exposure, but it is usually more time-consuming (Schubert et al. 2014). Prompted by the discovery of abnormal patterns of death in preliminary studies of vehicle control larvae housed in the same 96-well plate as DFP-exposed larvae, we hypothesized that DFP volatilizes and cross-contaminates wells in the 96-well plate exposure format. Our hypothesis was based on the fact that while DFP dissolves readily into phosphate-buffered saline for direct injection into mammalian subjects, its logKow of 1.13 and vapor pressure of 0.579 mm Hg at 20°C suggest that DFP volatilizes from water surfaces (NCBI 2021a; NCBI 2021b). To test this hypothesis, we conducted several survivability assays (comparing larvae exposed to all DFP concentrations on the same plate to control plates containing one exposure concentration per plate) and measured AChE specific activity in zebrafish exposed to DMSO and DFP on the same plates in comparison to DMSO-only larval controls exposed on separate plates.

## MATERIALS AND METHODS

### Zebrafish husbandry

Fish husbandry, spawning, and experiments were performed with the approval of the University of California, Davis (UC Davis), Institutional Animal Care and Use Committee. Fish husbandry and spawning were conducted according to previously described methods (Mundy et al. 2021). Tropical 5D wild type zebrafish were obtained from Sinnhuber Aquatic Research Laboratory (SARL) at Oregon State University, Corvallis, OR, and subsequent generations were raised at UC Davis. Spawning was conducted overnight in false bottom chambers, with embryos subsequently collected and transferred to plastic petri dishes containing embryo medium (EM) (Westerfield and ZFIN 2000). All experiments were conducted using 5 days postfertilization (dpf) larvae, and each experiment included replicates using embryos obtained from at least three separate spawning events.

### Chemical source information

DFP was purchased from Sigma-Aldrich (St. Louis, MO, USA) (∼90±7% purity), and stored as neat stock in -80°C. DFP was dissolved in either 100% DMSO (Sigma) or embryo media to create a 1,000x stock the day of use. Further dilution to 2x sub-stocks was conducted using embryo medium immediately before exposure. Vehicle was 0.1% DMSO for all exposure paradigms except the paradigm specifically including no vehicle carrier.

### Survivability assays

Zebrafish larvae were raised in petri dishes until 4 dpf. At 4 dpf, individual larvae were randomly chosen and transferred into a single well of a 12×8 96-well plate containing 50 µL EM, covered with Parafilm® to limit evaporation, and allowed to acclimate overnight in a 29°C incubator. The plates used for these experiments were either Falcon® polystyrene treated plates (REF 353075, Corning Inc., Corning, NY) (used for all experiments except for one specifically testing non-plasma-treated plates) or Costar polystyrene non-treated plates (REF 3370, Costar, Kennebunk, ME). At 5 dpf, larvae were exposed to DFP or vehicle via the addition of 50 µL of 2x sub-stock in embryo media. The plates were covered with Parafilm® and placed in 29°C incubator until removed for either larvae survivability assay or euthanization for AChE activity assay.

For all survivability assays except the control plates and randomized experiment, larvae were exposed to vehicle (0.1% DMSO) or varying concentrations of DFP ranging from 0.01 Mm – 1mM in the first six columns of the plate and repeated in columns 7-12 (Figure 1A and S1). In the randomized experiment, larvae were exposed to vehicle or 0.01 mM – 1 mM DFP in randomized groups of four across the entire plate (Figure S2). Control fish in a separate plate were exposed to DMSO-only or one concentration of DFP in two columns of the plate only (e.g., DMSO-only in column 1 and 7, 0.01 mM DFP in columns 2 and 8, etc.) (Figure 1A).

**Figure 1.**
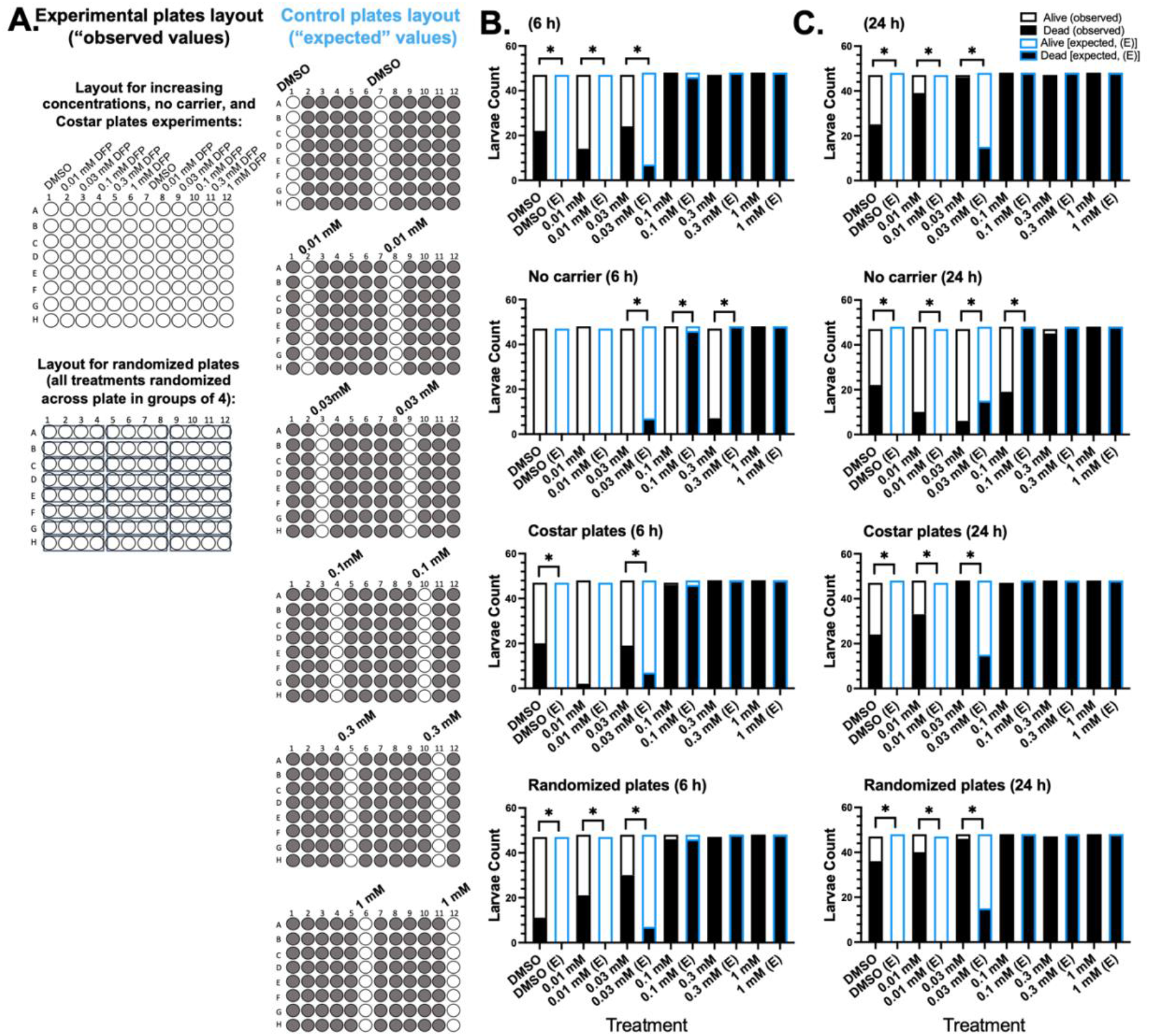
**A)** Diagrams of positional experimental design for experimental plates and control plates. All plates (experimental and control) were completed in triplicate. Grey circles represent wells not used. **B)** Mortality measured at 6 h of exposure. **C)** Mortality measured at 24 h of exposure. n=47-48 per group. * p < 0.05 in Fischer’s exact test, in comparison to expected values for each group.

At 6 h and 24 h, larvae were observed under a light microscope. Deceased larvae were counted and their position on the plate recorded. The conditions tested in the survivability assays included 1) increasing DFP concentrations across wells, 2) increasing DFP concentrations across wells using a brand of 96-well plate that is not treated with plasma-gas (Costar), 3) increasing DFP concentrations across wells using embryo media only as a carrier for DFP, and 4) randomizing the exposures across the plate. Larvae lacking a heartbeat were considered dead. Each experiment was conducted in triplicate (n=3 separate plates) using 16 larvae per exposure group (n = 48 larvae per experimental group).

### Acetylcholinesterase (AChE) activity assay

Larvae were exposed according to the exposure paradigm described above (“Survivability assays” section). Separate plates of control fish were exposed to vehicle in columns 1 and 7. After 1 h of exposure, larvae were removed from the wells and processed in pools of two, such that each column provided four samples of the same exposure group. In total, 56 samples were collected from each exposure plate, and 8 samples from each control plate (Figure 2A). The experiment was done in triplicate (n=3 larval pairs per spatial position).

**Figure 2.**
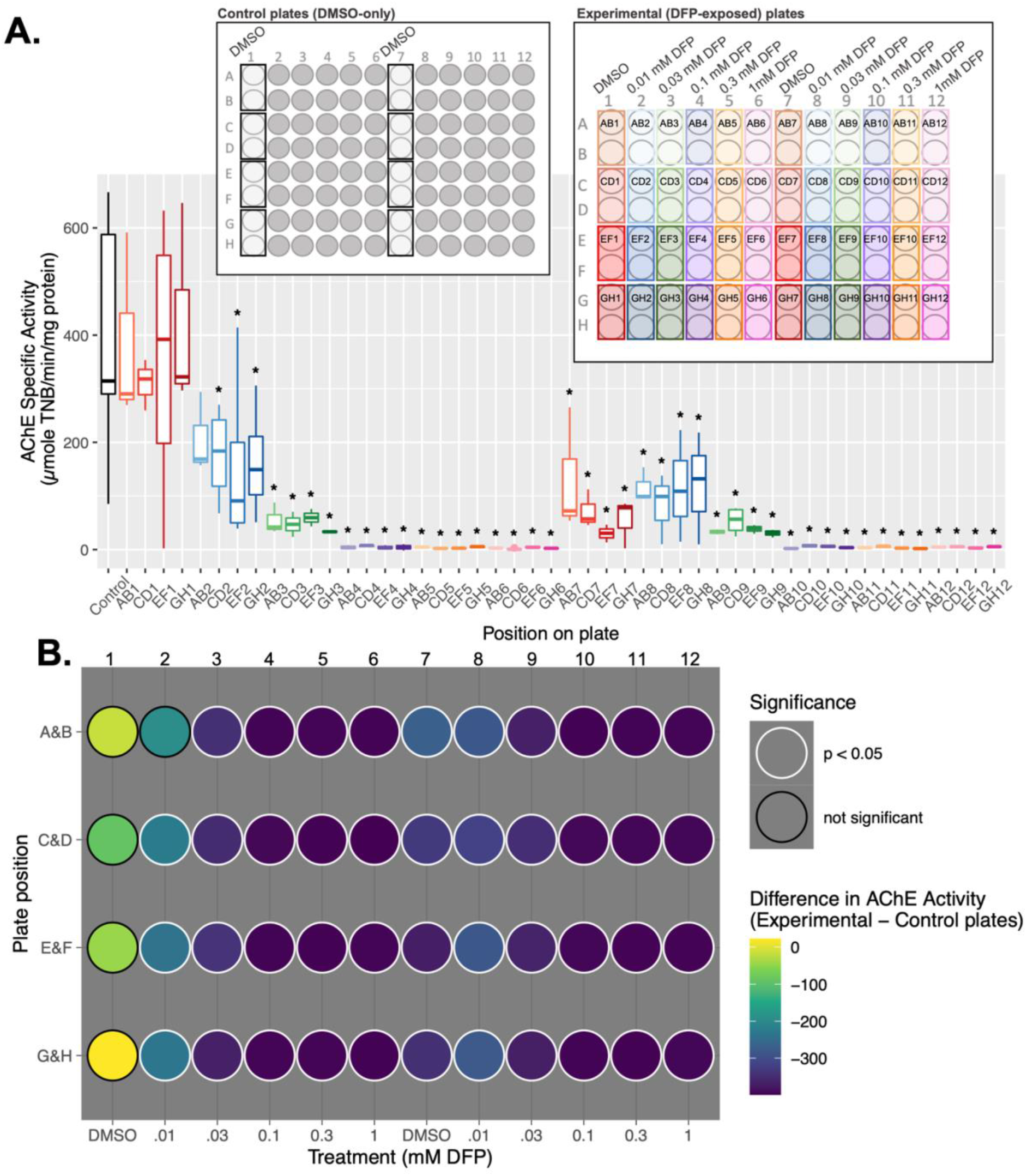
**A)** Boxplot represents AChE specific activity. Schematics show positional exposure paradigm. Boxes within the schematics represent pairs of larvae that were pooled for AChE activity assay, and labels within the boxes correspond to the labels on the x axis of the boxplot. n=3, * p < 0.05 in Dunnett’s multiple t test in comparison to control. **B)** Heatmap shows the difference of AChE activity in exposed plates subtracted from control plates. p < 0.05 in Dunnett’s multiple t-test in comparison to control (n=3).

The AChE activity assay was conducted as previously described in (Yang et al. 2011). The larvae were rinsed with DI water, euthanized on ice, and then homogenized in 120 µL of cold 1X phosphate-buffered saline (PBS) containing 1% TritonX-100 using a handheld homogenizer with disposable pestle. Lysates were snap-frozen in liquid nitrogen and stored at -80°C until use. Lysates were thawed and spun at 12,000 X g for 1 minute. The supernatant was added to a 96-well plate in triplicate, and AChE activity was measured using the standard Ellman assay (Ellman et al., 1961) with 5,5’-Dithiobis (2-nitrobenzoic acid) (DTNB) and acetylthiocholine iodide (ASChI) as substrates (Sigma-Aldrich). All plates contained three blanks with DTNB. Plates were equilibrated at room temperature for 5 min, and the reaction was initiated by the addition of ASChI. Absorbance at 405 nm was measured every two minutes for 30 minutes on a Synergy H1 hybrid reader (BioTek, Winooski, VT). AChE activity was normalized to total protein concentration of each sample as determined by a BCA assay (Pierce, Rockford, IL).

### Statistical analysis

#### Survivability assays

To specifically measure the effect of exposing larvae on the same plate in comparison to separate plates, the total dead larvae in the control plates (plates that contained only one exposure concentration) were considered the “expected” values (Figure 1A). Each “observed” value (the total dead larvae in the experimental plates) was compared to the corresponding concentration of DMSO or DFP on the control plates (the “expected” values) using two-sided Fischer’s exact test (α < 0.05). Survivability assay statistics were completed using GraphPad Prism (Version 9.1.0). All experimental plates including visual representation of spatial position of death at 6 and 24 h are shown in Figures S1 and S2.

#### AChE activity assay

To ensure the control plates did not have spatial differences in AChE activity, a non-parametric Kruskal Wallis ANOVA and post-hoc Dunn’s pairwise comparison were completed to compare each spatial position with the other. Because no significant differences were found (Figure S3), the values from all control samples (n=24) were combined and used as the control for comparison to each position on the exposed plates. Each spatial position (n=3) was compared to the control samples in non-parametric Kruskal Wallis ANOVA and post-hoc Dunnett’s multiple t-test (α < 0.05). AChE activity assay statistics were completed using R (version 4.0.3) (R Core Team 2021), and packages rstatix (Kassambara 2021) and DescTools (Signorell 2021).

## RESULTS

### Survivability assays

To investigate whether the cause of death in the vehicle control zebrafish observed in preliminary studies was caused by DFP cross-contamination, we first conducted survivability assays. The conditions tested in the survivability assays were 1) increasing DFP concentrations across wells, 2) increasing DFP concentrations across wells using a brand of 96-well plate that is not treated with plasma-gas (Costar brand), 3) increasing DFP concentrations across wells using embryo media only as a carrier for DFP, and 4) randomizing the exposure groups across the plate.

Larvae exposed to increasing concentrations of DFP on the same plate exhibited significantly more death (in the DMSO, 0.01 mM, and 0.03 mM DFP exposed wells) in comparison to control (larvae with similar exposures on separate plates) at 6 h and 24 h of exposure (Figure 1B, 1C). When the same experiment was conducted with no carrier (using DFP that was not prepared in DMSO), the larvae on the control plates (separated exposures) exhibited significantly more death at 6 h of exposure to 0.03, 0.1, and 0.3 mM DFP (Figure 1B). At 24 h of exposure, the larvae exposed on the same plate exhibited significantly more death compared to control plates in the vehicle and 0.01 mM DFP exposed zebrafish (Figure 1C). When using a different brand of plate (Costar), larvae exposed to increasing concentrations of DFP on the same plate exhibited significantly more death in vehicle DMSO and 0.03 mM DFP exposed wells at 6 h, and vehicle, 0.01, and 0.03 mM DFP exposed wells at 24 h in comparison to control plates (Figure 1B, 1C). Finally, when larvae were exposed to randomized concentrations of DFP on the same plate, significantly more death was observed in the vehicle, 0.01 mM, and 0.03 mM DFP wells at 6 h and 24 h of exposure in comparison to control plates (Figure 1B, 1C).

### AChE activity assays

To evaluate whether zebrafish were exposed to DFP, AChE activity was measured in zebrafish tissue in experimental plates in which larvae were exposed to vehicle and increasing concentrations of DFP on the same plate for 1 h in comparison to separate control plates containing vehicle-only exposed larvae. Each position on the experimental plate was compared to the control values. AChE activity was found to be significantly decreased in all wells except the first column of vehicle control larvae and the top two wells of 0.01 mM DFP-exposed larvae. Most notably, all vehicle control larvae in column 7 (in the middle of the plate) exhibited significantly decreased AChE activity.

## DISCUSSION

Zebrafish are a powerful tool for toxicological screening (Tal et al. 2020) and are a desirable model for investigating individual toxic compounds such as DFP. However, a limitation of using zebrafish larvae in a medium-high throughput screen involves dosing the animals via their aqueous environment rather than direction injection. Here, we investigated whether the common 96-well plate exposure method is appropriate for testing larval zebrafish exposure to semi-volatile chemicals, such as DFP.

In the survivability assays, the most notable observation was the consistent and significant death of the vehicle control larvae on experimental plates that also contained wells of zebrafish exposed to increasing concentrations of DFP (Figure 1A, 1B). Further, the vehicle control larvae on the experimental plates spatially positioned in the middle of the plate (column #7) showed more and earlier (6 h as opposed to 24 h) death compared to the vehicle control larvae in the leftmost column (column #1) (Figure S1) that were furthest away from the DFP containing wells.

To test whether the pretreatment (plasma gas coating) of the polystyrene Falcon Corning plates affected survivability, we repeated the experiment using non-treated polystyrene Costar plates. Polystyrene tissue culture plates are typically pretreated with plasma gas to deposit a negative charge at the bottom of the wells to enhance cell attachment. To the best of our knowledge, the pretreatment of culture plates with plasma gas has not been found to have negative impacts on zebrafish larvae survival. Here we tested whether the pretreatment of polystyrene Falcon Corning plates is a confounding factor for DFP volatility or toxicity by repeating the experiment (increasing concentrations of DFP on the same plate) with non-treated polystyrene 96-well plates (Costar brand). Because the results were almost identical to the Falcon Corning plates (Figure 1A, 1B), we concluded that plasma gas pretreatment of the 96-well plate is not an important factor in whether DFP volatilizes and cross-contaminates wells.

In order to test whether exposure position on the plate had an effect on death, we positioned vehicle or DFP exposures in a randomized fashion (Figure 1A, Figure S2). The results were similar to the experimental design in which larvae were exposed to increasing concentrations of DFP across the plate in that the vehicle control, 0.01, and 0.03 mM DFP-exposed larvae exhibited significantly increased death compared to control plates (separate exposures) at 6 and 24 h of exposure (Figure 1B, 1C). These results indicate that the spatial positioning on the plate alone does not influence the rate of death; however, the proximity to high concentrations of DFP may influence the rate of death.

Interestingly, the larvae exposed to DFP without carrier (DFP was dissolved in embryo media directly without DMSO) exhibited significantly lower death than larvae exposed on separate plates to 0.03, 0.1, and 0.3 mM DFP for 6 h, and 0.03 and 0.1 mM DFP for 24 h. However, at 24 h of exposure, vehicle and 0.01 mM DFP-exposed larvae on the experimental plates exhibited significantly more death compared to control plates. These results suggest that 1) DMSO increases the volatilization of DFP, as evidenced by decreased amounts of cross contamination occurring within 6 h of exposure when DMSO is not used as a vehicle, 2) DFP may not be fully soluble in embryo media and coming out of solution, and 3) the presence of DMSO facilitates better delivery to zebrafish tissues.

In an AChE assay conducted to confirm the distribution of DFP across the plate, vehicle control larvae had significant AChE activity inhibition. Thus, we confidently conclude that DFP can volatilize and cross-contaminate wells within the same plate (Figure 2A, 2B). Notably, the AChE assay was conducted at 1 h after exposure, and no death was observed in any experimental group at this time point.

Temperature and vehicle may be two factors that play an important role in the volatility of DFP. Considering the design of all experiments presented here, it is important to note that all tests were conducted at 29°C, and all experiments (excluding the tests specifically using no carrier) used DMSO as the vehicle carrier. It may be worth exploring the possibility of exposing zebrafish to DFP using the 96-well plate method under different conditions (i.e., using a lower temperature or different vehicle such as methanol or acetone) in order to maintain the high-throughput ability of the assay while minimizing cross contamination.

## CONCLUSIONS

In summary, our data support the hypothesis that DFP volatilizes and cross-contaminates wells when placed in embryo media in a 96-well plate with lid and Parafilm® at 29°C using DMSO as a vehicle. This observation is important when considering experimental design for studying the effects of DFP on zebrafish larvae. Based on the findings of this study, we strongly support the use of different exposure methods when testing the effects of semi-volatile and volatile reagents on zebrafish larvae (i.e., exposing zebrafish larvae using individual cell culture plates, separating exposure groups on individual 96-well plates as opposed to exposing larvae to all concentrations on one plate, using a separate plate for vehicle control, or microinjection). Lastly, these observations serve to inform other investigators of the possibility of cross-contamination with compounds of similar chemical properties to DFP when using the 96-well plate exposure method.

## Acknowledgment

We gratefully acknowledge Donald Bruun and Cris Grodzki for consulting on the acetylcholinesterase inhibition assays. We also thank Dr. Suzette Smiley-Jewell (UC Davis CounterACT Center) and Dr. Heike Wulff (UC Davis Dept. of Pharmacology) for reviewing and editing this manuscript. This work was supported by the National Institute of Neurological Disorders and Stroke CounterACT program [grant number U54 NS079202 to PJL]. The sponsors were not involved in the study design, the collection, analysis, and interpretation of data, in the writing of the report or in the decision to submit the paper for publication.

