## Supplementary material for "Diisopropylfluorophosphate (DFP) volatizes and cross-contaminates wells in a common 96-well plate zebrafish larvae exposure method"

**SUPPLEMENTARY FIGURES**

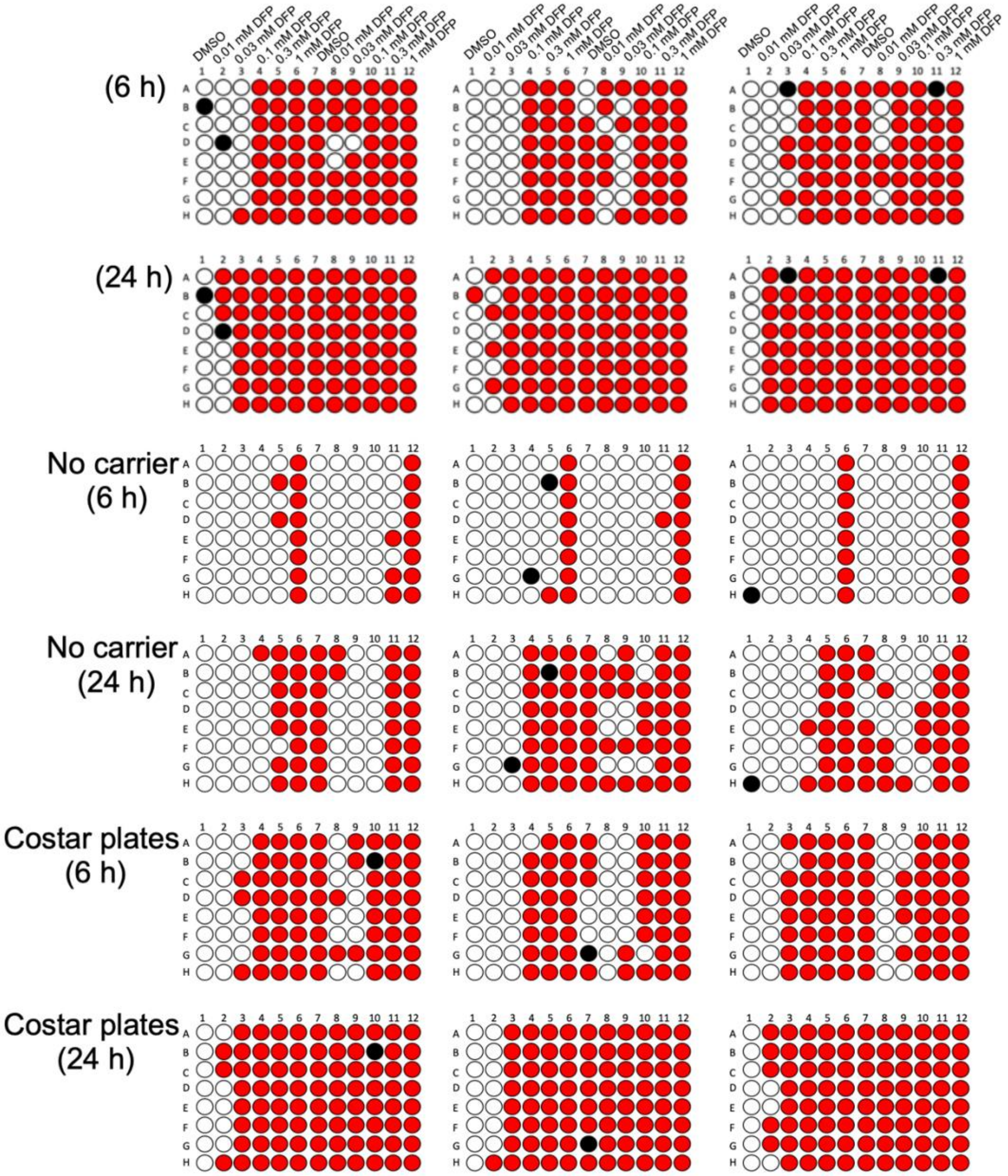


**Figure S1.** All individual experimental plates are shown. Each plate is one replicate, and each circle is one well containing an individual larva. Red wells represent deceased larvae, white wells represent live larvae at the time of observation, and black wells represent wells that were not counted (e.g., larvae were dead or deformed before exposure).

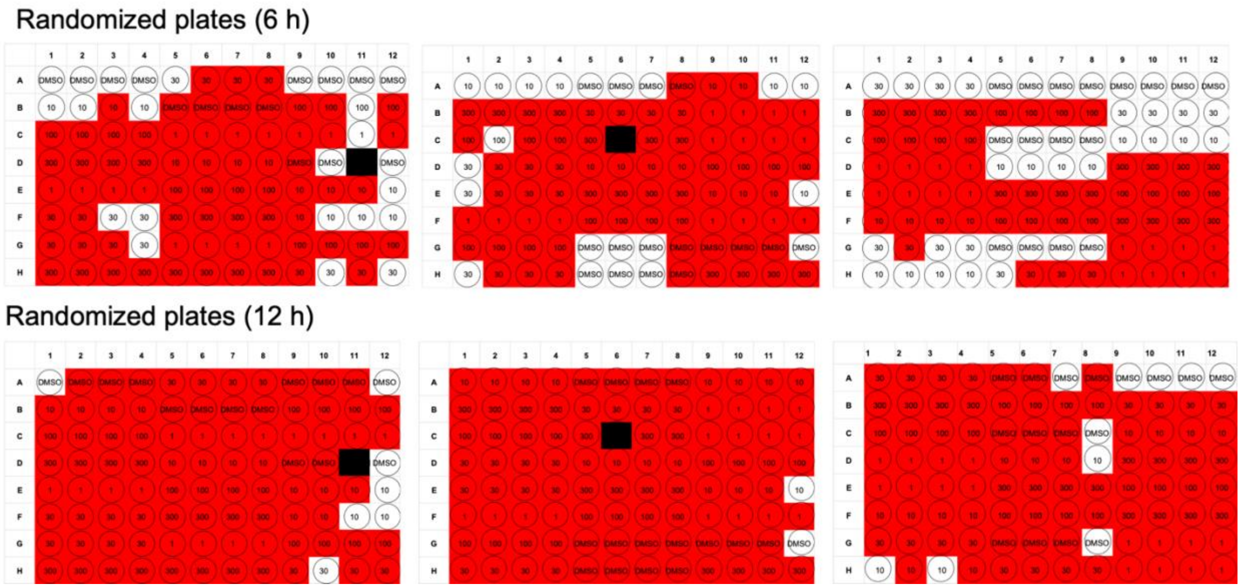


**Figure S2.** All individual experimental randomized plates are shown. Each plate is one replicate, and each circle is one well containing an individual larva. Red wells represent deceased larvae, white wells represent live larvae at the time of observation, and black wells represent wells that were not counted (e.g. larvae were dead or deformed before exposure).

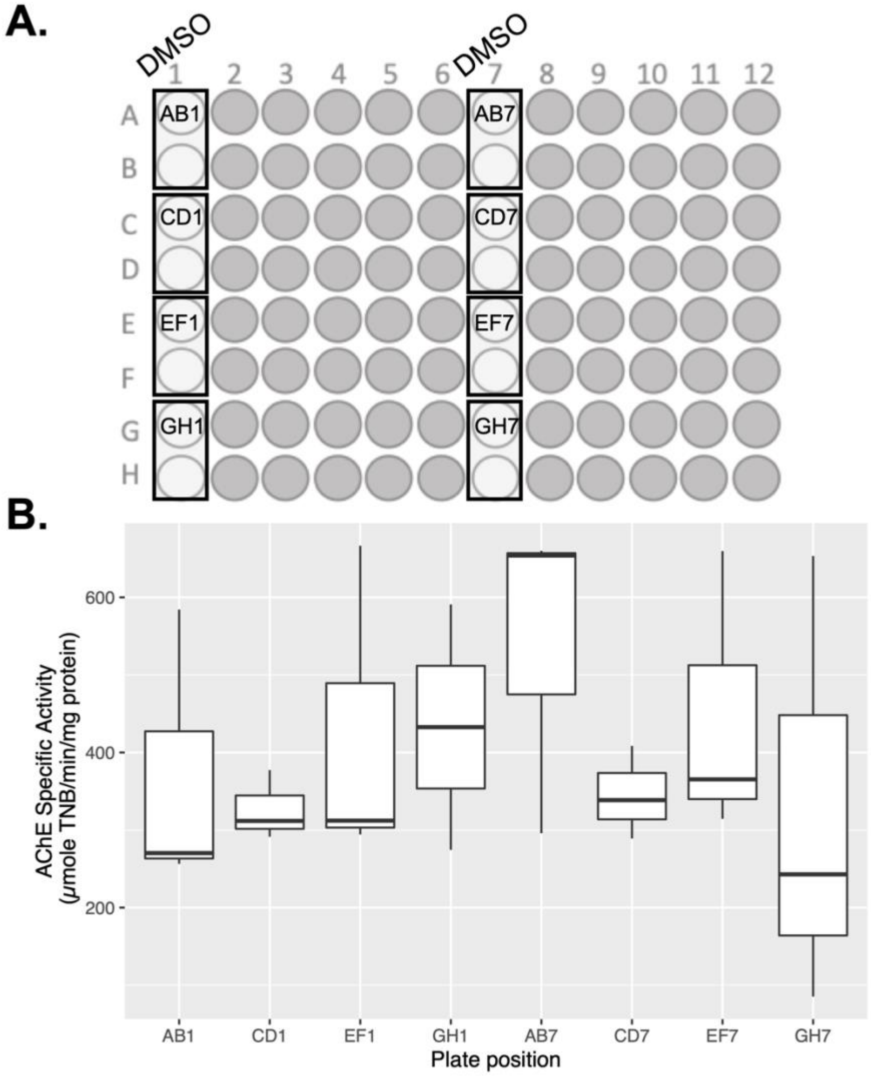


**Figure S3.** AChE activity spatially across control plates. **A)** Schematic showing the placement of DMSO-exposed larvae on the plates. Squares represent wells that were collected together for processing. Dark grey circles represent wells that did not contain zebrafish larvae. **B)** Boxplot showing the comparison of AChE activity across plate positioning. No significant differences were found between plate positions. AChE activity was compared using Dunn's multiple t-test ($\alpha < 0.05$), in which each position was compared against each other. (n=3).
